# Effects of gestational age at birth on perinatal structural brain development in healthy term-born babies

**DOI:** 10.1101/2021.05.02.442327

**Authors:** Oliver Gale-Grant, Sunniva Fenn-Moltu, Lucas França, Ralica Dimitrova, Daan Christaens, Lucilio Cordero-Grande, Andrew Chew, Shona Falconer, Nicholas Harper, Anthony N Price, Jana Hutter, Emer Hughes, Jonathan O’Muircheartaigh, Mary Rutherford, Serena J Counsell, Daniel Rueckert, Chiara Nosarti, Joseph V Hajnal, Grainne McAlonan, Tomoki Arichi, A David Edwards, Dafnis Batalle

## Abstract

Multiple studies have demonstrated less favourable childhood outcomes in infants born in early term (37-38 weeks gestation) compared to those born at full term (40-41 weeks gestation). While this could be due to higher perinatal morbidity, gestational age at birth may also have a direct effect on the brain and subsequent neurodevelopment in term-born babies. Here we characterise structural brain correlates of gestational age at birth in healthy term-born neonates and their relationship to later neurodevelopmental outcome. We used T2 and diffusion weighted Magnetic Resonance Images acquired in the neonatal period from a cohort (n=454) of healthy babies born at term age (>37 weeks gestation) and scanned between 1 and 41 days after birth. Images were analysed using tensor based morphometry (TBM) and tract based spatial statistics (TBSS). Neurodevelopment was subsequently assessed at age 18 months using the Bayley-III Scales of Infant and Toddler Development, and the effects of gestational age at birth and related neuroimaging findings on outcome were analysed with linear regression. Infants born earlier had areas of higher relative ventricular volume, and lower relative brain volume in the basal ganglia, cerebellum and brainstem. Earlier birth was also associated with lower fractional anisotropy, higher mean, axial and radial diffusivity in major white matter tracts. Gestational age at birth was positively associated with all Bayley-III subscales at age 18 months. Linear regression models predicting outcome from gestational age at birth were significantly improved by adding neuroimaging features associated with gestational age at birth. This work adds to the growing body of evidence of the impact of early term birth and highlights the importance of considering the effect of gestational age at birth in future neuroimaging studies including term-born babies.

## Introduction

Study of inter-individual variation in the duration of pregnancy has taken place for at least 2500 years (Malamitsi-Puchner, 2017). Academic consensus for the vast majority of this time has been that childbirth after 37 weeks gestational age (usually defined as the time since the first day of the mother’s last menstrual period) is optimal, and infants born earlier than this are at an increased risk of negative outcomes (Budin, 1907). Recent research has challenged the idea of a term - preterm binary classification and proposed that babies born earlier within the 37 to 41-week (term) range have less desirable outcomes than those born later. The first large study to demonstrate this, published in 2012, used a cohort of 18,818 infants and found that individuals born between 37 and 38 weeks gestational age had a higher risk of adverse health and developmental outcomes at age 3 compared to those born between 39 and 41 weeks gestational age (Boyle et al., 2012). Subsequent studies have linked other phenotypes with early term birth, including higher risks of neonatal (Murzakanova et al., 2020) and later childhood illness (Coathup et al., 2020) and less favourable neurodevelopmental outcomes in early childhood (Hua et al., 2019; Rose et al., 2013).

The underlying mechanism of these differences is not fully understood, although several hypotheses have been proposed. Brain development occurs rapidly in the third trimester – being born (and hence removed from the supportive maternal environment) earlier in this period may disrupt normal neurodevelopment (Davis and Sandman, 2010). Outcome differences could also be due to perinatal risk factors: infants born earlier have lower birthweights and a higher risk of perinatal morbidity (Murzakanova et al., 2020), which may lead to worse childhood neurodevelopmental outcomes (Doctor et al., 2001; Taylor et al., 2004). In many countries mothers who experience complications during pregnancy which themselves may adversely affect the baby, such as pre-eclampsia or intrauterine growth restriction, often have labour medically induced at a gestational age of 37 weeks, which could also contribute to differences in perinatal risk and on average poorer outcomes in early term born infants (Coates et al., 2020).

A small number of studies have examined imaging correlates of gestational age at birth in term born babies, all exclusively using diffusion tensor imaging acquired in the early neonatal period (Broekman et al., 2014; Jin et al., 2019; Ou et al., 2017). These studies all report similar results, namely lower fractional anisotropy (FA) and higher mean diffusivity (MD) in infants born earlier in the term period. Associations of gestational age at birth (GAAB) with MRI features in term only populations have also been reported in later childhood, including lower total brain volume in a cohort of 10 year old children (El Marroun et al., 2020), and a positive association of GAAB with functional connectivity in a cohort of 6-10 year olds (Kim et al., 2014).

One likely reason for the paucity of studies demonstrating a brain effect of GAAB in the neonatal period is that the majority of neonatal MRI studies thus far have focused on preterm or clinical risk populations (Batalle et al., 2018). However, this limitation can be overcome with the developing Human Connectome Project (dHCP) data which consists of high quality neonatal MRI data and complementary neurodevelopmental outcomes at age 18 months from a large cohort of healthy term-born individuals (Hughes et al., 2017), allowing us to test the hypothesis that GAAB will be correlated with neonatal brain structure in term-born babies (independently of age at scan). To do so, we analysed the volumetric (using tensor based morphometry) and white matter (using tract based spatial statistics) brain correlates of GAAB and their association to neurodevelopmental outcomes in a sample of 454 term-born neonates.

## Methods

### Sample

This study is based on a prospective sample of neonates participating in the Developing Human Connectome Project (dHCP, http://www.developingconnectome.org/). This project has received UK NHS research ethics committee approval (14/LO/1169, IRAS 138070), and written informed consent was obtained from parents.

For the purposes of this study, we have included only healthy term-born singletons. Exclusion criteria were as follows: GAAB < 37 weeks, admission to neonatal intensive care following birth, non-singleton birth, medical complication reported during pregnancy by mother, intrauterine growth restriction or small for gestational age birthweight, or a visible abnormality of possible clinical or analytical significance on MRI following reporting by an experienced Paediatric Neuroradiologist.

### Image Acquisition

Neonatal MRI data were acquired on a Phillips 3 Tesla Achieva system (Philips Medical Systems, Best, The Netherlands) at the Evelina Newborn Imaging Centre, Evelina London Children’s Hospital. All infants were scanned during natural sleep without sedation as previously described (Hughes et al., 2017), including a bespoke transport system, positioning device and an optimally sized neonatal 32-channel receive coil with a custom-made acoustic hood. All scans were supervised by a neonatal nurse and/or paediatrician who monitored heart rate, oxygen saturation and temperature throughout the scan.

T2-weighted images were obtained using a Turbo Spin Echo (TSE) sequence, acquired in two stacks of 2D slices (in sagittal and axial planes), using parameters: TR=12s, TE=156ms, SENSE factor 2.11 (axial) and 2.58 (sagittal), acquisition resolution 0.8×0.8×1.6mm^3^ with 0.8mm slice overlap, reconstructed to 0.8 mm^3^ isotropic resolution, and diffusion images were acquired using parameters TR=3800ms, TE=90ms, SENSE factor=1.2, multiband factor=4, acquisition resolution 1.5×1.5×3.0mm^3^ with 1.5mm slice overlap. Diffusion gradient encoding included images collected at b=0s/mm^2^ (20 repeats), b=400s/mm^2^ (64 directions), b=1000s/mm^2^ (88 directions), b=2600s/mm^2^ (128 directions) (Hutter et al., 2018), and images were reconstructed to a final resolution of 1.5×1.5×1.5mm^3^ (Christiaens et al., 2021; Hutter et al., 2018).

### Image Processing

Motion-correction and slice-to-volume reconstruction of T2 weighted images was carried out using a dedicated algorithm as previously described (Cordero-Grande et al., 2018). These images were subsequently registered to a dHCP week-specific template (according to each individual’s postmenstrual age at scan) in a common space corresponding to gestational age of 40 weeks (Schuh et al., 2018) using the Symmetric Normalization (SyN) algorithm from Advanced Normalization Tools (ANTs), version 3.0 (Avants and Gee, 2004). Affine and non-linear transformations were performed, with the last of these used to create deformation tensor fields in template space to remove the effect of individual variations in head size from the final analysis. The resulting tensor fields were used to calculate scalar Jacobian determinants, which were subject to logarithmic transformation, using ANTs. Log-Jacobian determinant maps (hitherto referred to as Jacobians) were smoothed with a sigma of 3.5 mm full width at half maximum (FWHM) Gaussian filter. Images were finally resized to an isotropic voxel size of 1mm prior to Tensor Based Morphometry (TBM) analysis (Ashburner and Friston, 2000).

Diffusion MRI was pre-processed as previously described (Kelly et al., 2019). Briefly, images were denoised (Cordero-Grande et al., 2018), Gibbs ringing suppressed (Kellner et al., 2016) and reconstructed using a slice-to-volume motion correction technique that uses a bespoke spherical harmonics and radial decomposition (SHARD) method, together with outlier rejection, distortion and slice profile correction (Christiaens et al., 2021). Tensors were reconstructed and non-linearly registered to a population-based template using DTI-TK (Zhang et al., 2006). Mean diffusivity (MD), axial diffusivity (AD), radial diffusivity (RD) and fractional anisotropy (FA) maps for each subject were subsequently generated in template space, and re-integrated into the FSL tract based spatial statistics (TBSS) pipeline optimised for the neonatal brain (Ball et al., 2010; Jenkinson et al., 2012).

### Follow-up

Neurodevelopmental assessment was offered to all participants at 18 months of age. Neurodevelopmental performance was assessed using the Bayley Scales of Infant and Toddler Development, Third Edition (Bayley, 2006). The cognitive, motor and communication composite scores are used for analyses in this study. Index of multiple deprivation (IMD), (a geographically defined composite social risk score comprising data on income, employment, health, education, living environment and crime) was calculated from the mother’s home address at the time of birth and included as a covariate in all associations with follow-up outcome data (Abel et al., 2016).

### Statistical Analysis

TBM and TBSS analyses were performed using the *randomise* tool for nonparametric permutation inference in FSL with 10,000 permutations per test (Winkler et al., 2014). Threshold free cluster enhancement and family wise error (FWE) rate was applied to correct for voxel-wise multiple comparisons. All general linear models (GLM) contained post menstrual age at scan and sex as covariates.

Associations between GAAB, imaging phenotypes and Bayley-III scores were performed using univariate linear regression in SPSS statistics 26. Graphical figures are drawn in GraphPad Prism 8.

Specifics of tests used for associations of GAAB and neurodevelopmental outcome are indicated with each result.

### Data availability

The dHCP is an open-access project. The imaging and demographic data used in this study can be downloaded by registering at https://data.developingconnectome.org/.

## Results

### Population

At the time of the study commencing, MRI data had been collected and pre-processed from 771 individuals. 251 individuals were excluded due to being born preterm, 12 due to nonsingleton birth, 18 due to medical complications of pregnancy, 26 due to a birthweight < 10^th^ centile (small for gestational age) and 10 due to visible abnormalities of possible clinical significance on MRI. This left 454 individuals included in the study. Of the 454 individuals, 374 had diffusion MRI data, 332 had T2 data and were followed up at 18 months, and 281 had T2 data, DWI data, and were followed up at 18 months. The demographic details of the group are summarised in Table 1.

**Table 1 –.** Demographic details of cohort.

|  | T2 MRI | Diffusion MRI | Follow-up and T2 MRI | Follow-up & both MRI modalities |
| --- | --- | --- | --- | --- |
| **N** | 454 | 374 | 332 | 281 |
| **Male** | 247 | 204 | 179 | 146 |
| **Female** | 207 | 170 | 153 | 135 |
| **Gestational age at birth [weeks]** | 40.1 (39.1 - 41.0) | 40.1 (39.3 – 40.7) | 40.1 (39.1 – 40.9) | 40.1 (39.4 – 41.0) |
| **Post menstrual age at scan [weeks]** | 41.3 (40.3 - 42.6) | 40.9 (40.3 – 42.0) | 41.2 (40.1 - 42.4) | 41.0 (40.4 – 42.1) |
| **APGAR (5 minute)** | 9 (8-10) | 9 (7-10) | 9 (8-10) | 9 (8-10) |
| **Birthweight Centile** | 52 (28-77) | 51 (29-76) | 57 (39-79) | 51 (29-77) |

Median (IQR) indicated.

### Brain Volume (TBM)

Voxel-wise analysis of relative brain volume using TBM showed a positive correlation of GAAB (corrected for postmenstrual age at scan and sex) with relative brain volume (Jacobians) of the deep grey matter, basal ganglia, medial cerebellum, parts of the inferior temporal lobes and parts of the brainstem bilaterally. The relative volumes of parts of the ventricles were negatively correlated with GAAB (Figure 1).

**Figure 1 –.**
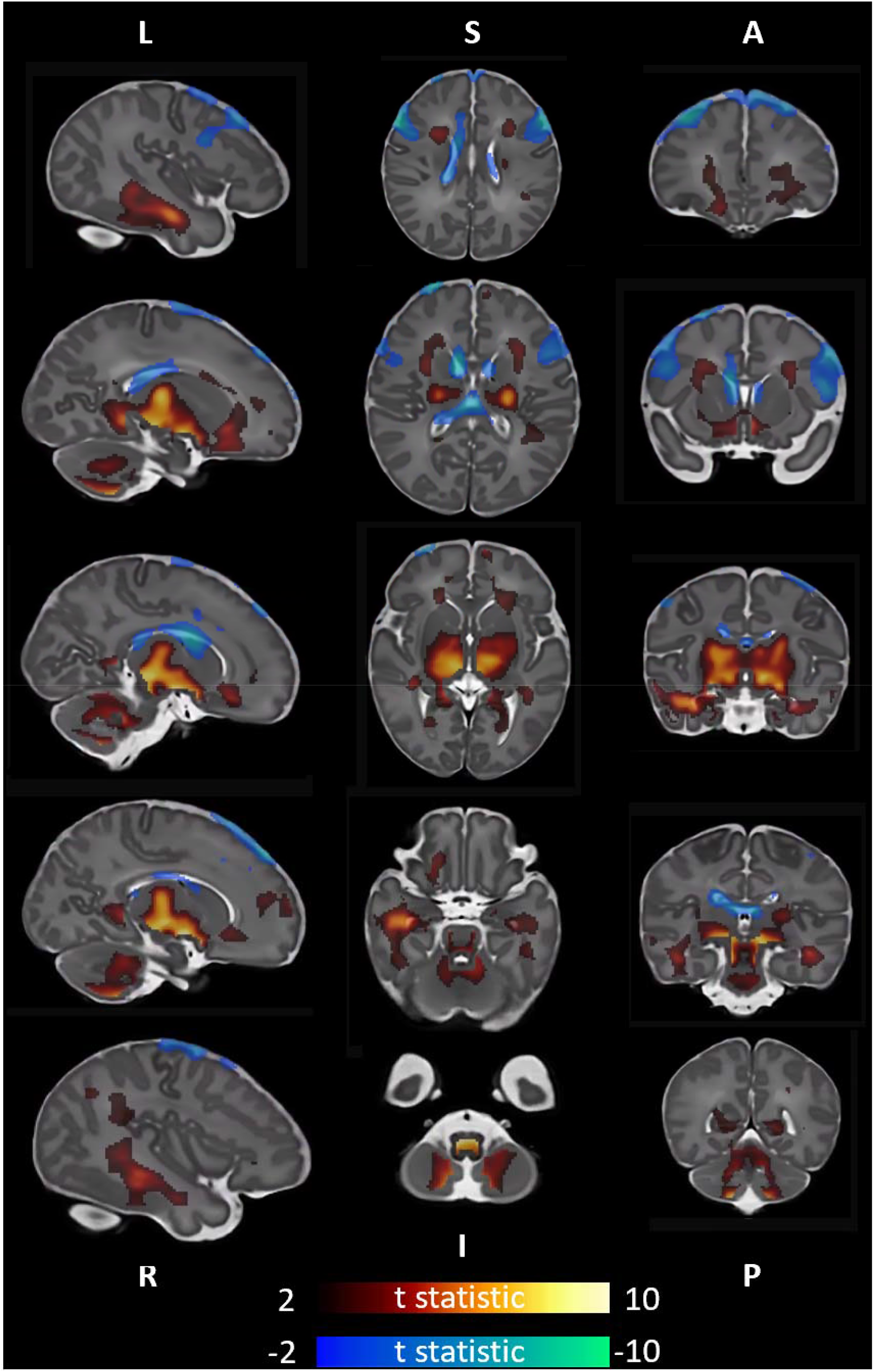
Association of gestational age at birth with brain volume. T statistic of areas of significant (p<0.025) correlation shown - positive in red-yellow, negative in blue-green. 1^st^ column left to right, 2^nd^ column superior to inferior, 3^rd^ column anterior to posterior. Analysis corrected for postmenstrual age at scan and sex.

### White Matter Diffusion Measures (TBSS)

Analysis of white matter diffusion MRI characteristics using TBSS showed that gestational age at birth was positively correlated with FA and lower MD, AD and RD. Associations with GAAB were apparent in multiple major white matter tracts but were most pronounced in the corticospinal tracts. Associations were most pronounced in axial diffusivity, and least pronounced in fractional anisotropy (Figure 2).

**Figure 2 –.**
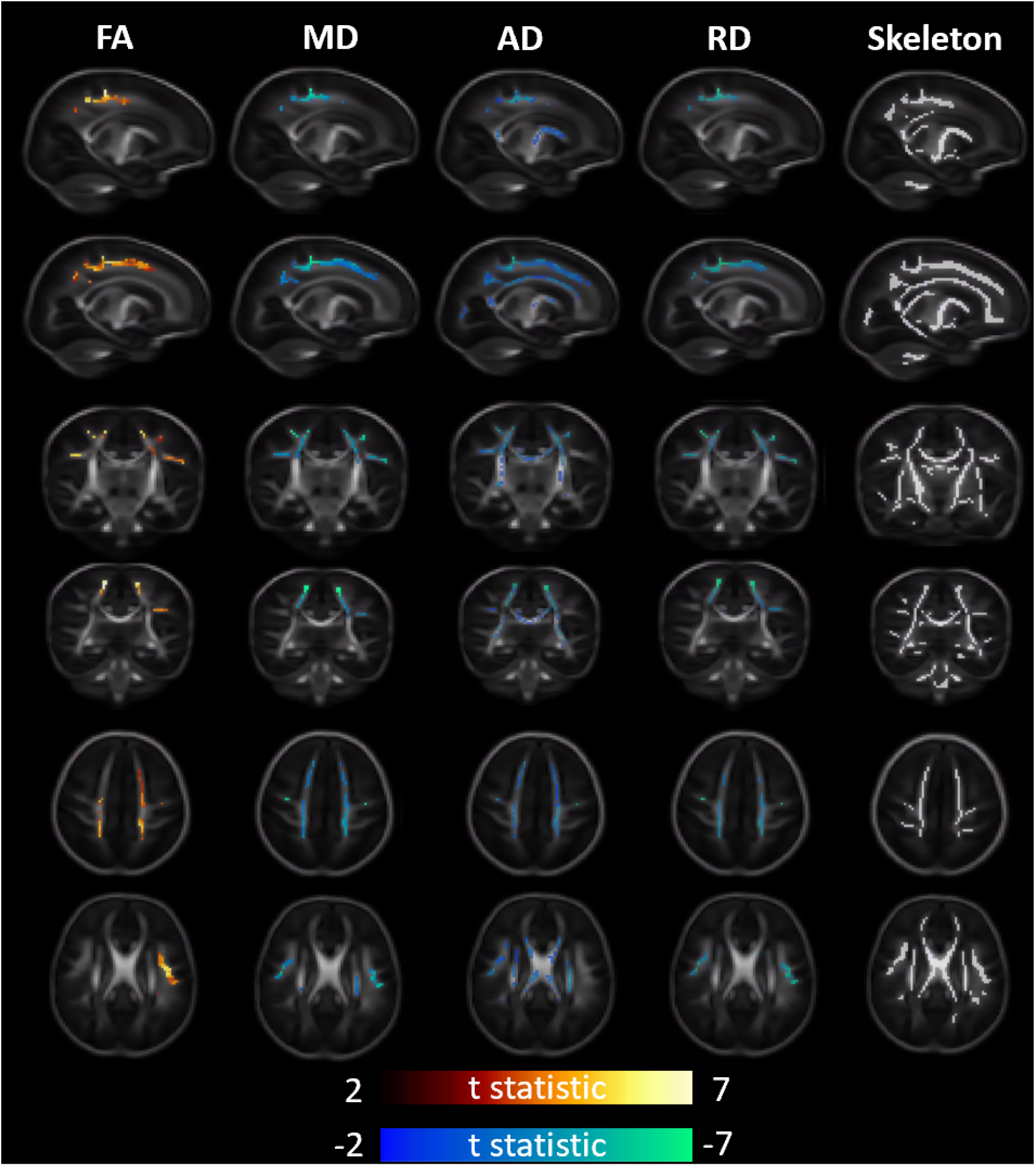
Association of gestational age at birth with white matter diffusivity measures. T statistic of areas of significant (p<0.025) correlation shown - positive in red-yellow, negative in blue-green. From left to right – Fractional Anisotropy, Mean Diffusivity, Axial Diffusivity, Radial Diffusivity, representative diffusion skeleton. From top to bottom – Axial (superior), Axial (inferior), Sagittal (L), Sagittal (R), Coronal (Anterior), Coronal (Posterior). Analysis corrected for post menstrual age at scan and sex.

### Neurodevelopmental Phenotype at 18 Months

In keeping with previous studies GAAB was positively correlated with all sub-domains of the Bayley-III Scales (Cognitive adj. r^2^ =0.04, p=0.002, Motor adj. r^2^ =0.04, p=0.001, Communication adj. r^2^ =0.04, p=0.001) (Figure 3).

**Figure 3 –.**
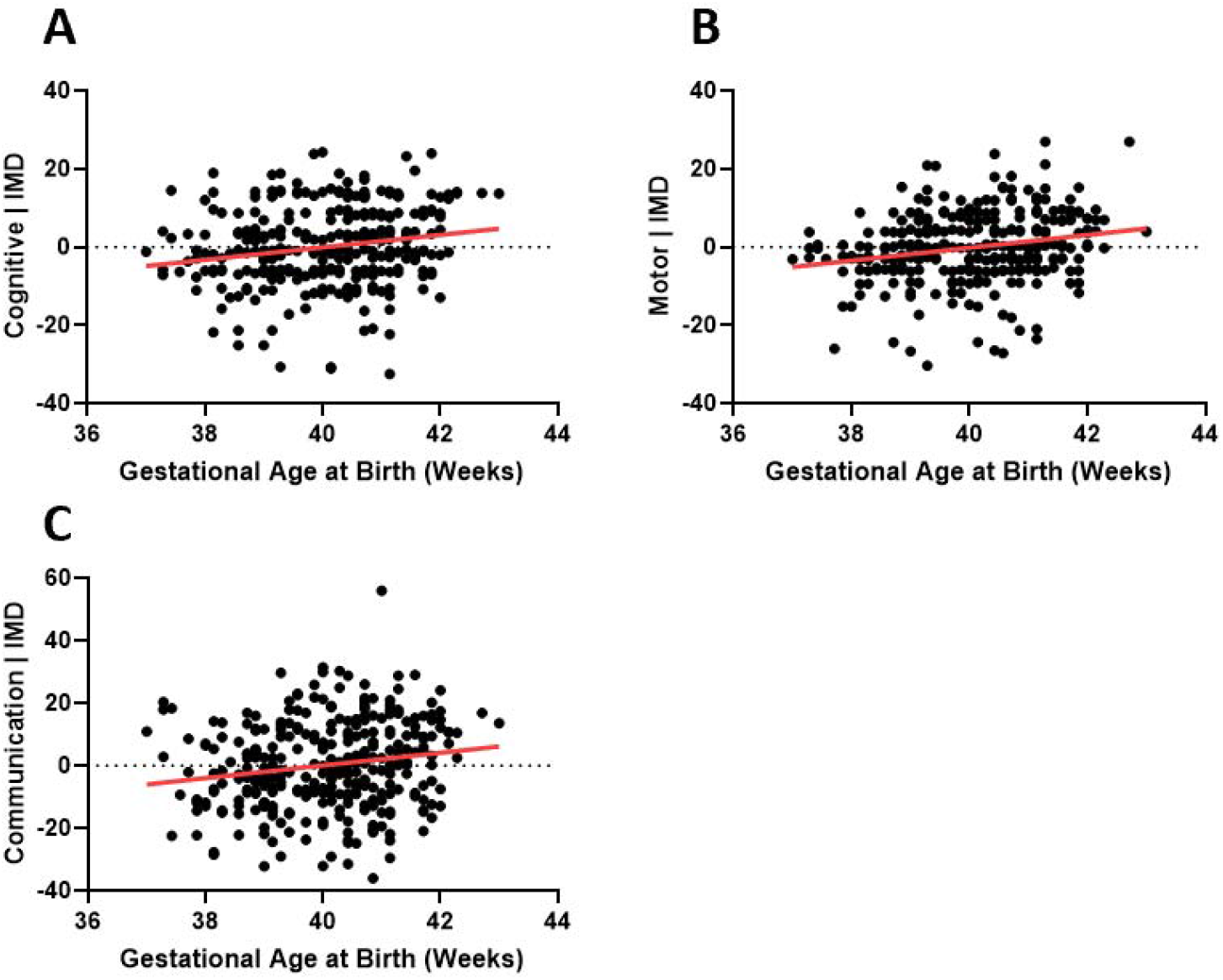
Association of gestational age at birth with developmental phenotype at 18 months (n=332). **A-C** Bayley-III Scales of Infant and Toddler Development **(A** Cognitive adj. r^2^ =0.04, p=0.002, **B** Motor adj. r^2^ =0.04, p=0.001, **c** Communication adj. r^2^ =0.04, p=0.001). All analyses corrected for Index of Multiple Deprivation score.

An exploratory analysis of the influence of the inter individual brain differences shown in Figures 1 and 2 on individual outcome differences was subsequently performed. For each neuroimaging modality the mean feature value of the region significantly associated with gestational age at birth was extracted. This gave six feature values per subject (volume positively associated with GAAB, volume negatively associated with GAAB, FA, MD, AD and RD associated with GAAB). The association of each of these values with the Bayley-III neurodevelopmental subscales was assessed using linear regression. As there are three subscales (Cognitive, Motor, Communication) this meant that 18 total analyses were performed, of which 2 were significant, although only one (volume negatively associated with GAAB – Bayley-III Motor Composite Score) would remain significant after correction for multiple comparisons (Figure 4).

**Figure 4 –.**
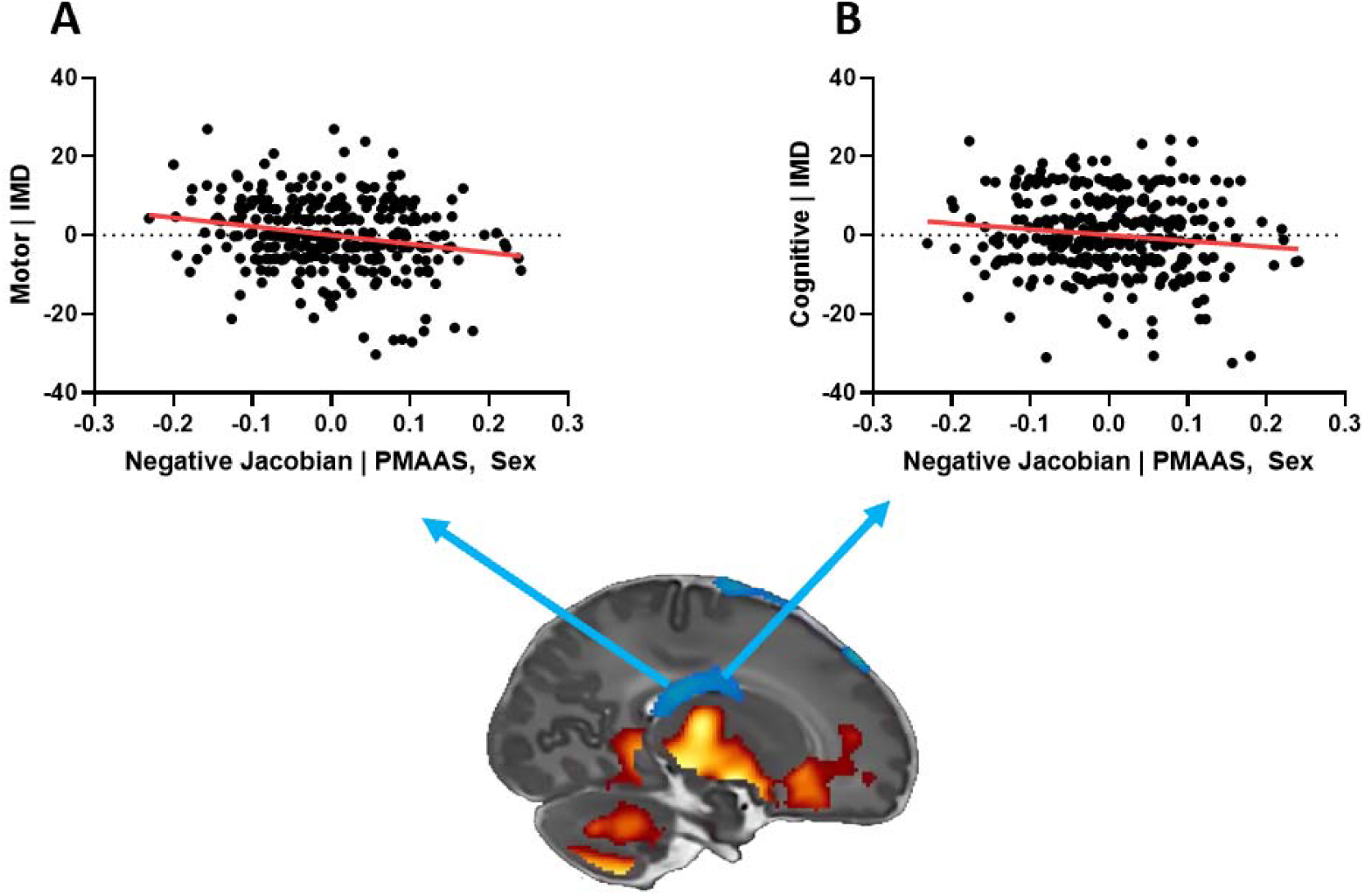
Association of regional volumes with developmental phenotype at 18 months (n=332). Regional volume negatively correlated with gestational age at birth associated with A Bayley-III Motor Score (adj. r^2^=0.04, p<0.001) and **B** Bayley-III Cognitive Score (adj. r^2^=0.02, p=0.02). Coloured arrows on sagittal section indicate region used in each analysis. Bayley-III scales corrected for Index of Multiple Deprivation score, mean regional volume feature values corrected for post menstrual age at scan and sex.

Next the mean feature values were added to each of the linear regression analysis of GAAB and Bayley-III outcome shown in Figure 4 (along with the neuroimaging covariates post menstrual age at scan and sex). Feature values were added in two groups – volume features and diffusion features. Analyses of the effect of adding brain volume information were performed in individuals who had a T2 scan and follow-up (n=332), and analyses of adding brain volume and white matter diffusion information were performed in individuals who had a T2 scan, a DWI scan and follow-up (n=281). Adding volume features significantly improved all models, whereas adding diffusion features only improved the Communication model (Table 2).

**Table 2 –.** Modulation of gestational age at birth - outcome association with neuroimaging findings. Analyses performed using univariate linear regression with separate models for Cognitive (A), Motor (B) and Communication (C) Composite Scores from the Bayley-III Scales. The basic model compares Gestational age at Birth and outcome with Index of Multiple Deprivation as a covariate. The Volume model adds the mean feature value (Jacobians) of regions positively or negatively associated with gestational age at birth, as well as the imaging covariates post menstrual age at scan and sex, and the DTI model adds the mean feature value of FA, MD, AD and RD regions associated with gestational age at birth, as well as the imaging covariates post menstrual age at scan and sex. Analysis of the effect of the Volume model assessed in all individuals who had a T2 scan and followup (n=332), analysis of the effect of the DTI and DTI + Volume models additionally assessed in individuals who had a T2 scan, a DWI scan, and follow-up (n=281).

| | df | Adjusted $r^2$ | Model p-value | F Change from GAAB + IMD | F Change p-value |
| --- | --- | --- | --- | --- | --- |
| **A. Cognitive** |  |  |  |  |  |
| **n=332** |  |  |  |  |  |
| Y = GAAB + IMD | 2 | 0.038 | 0.002 | - | - |
| Y = GAAB + IMD + [Volume] | 6 | 0.063 | 0.001 | 2.82 | 0.025 |
| **n=281** |  |  |  |  |  |
| Y = GAAB + IMD | 2 | 0.046 | 0.001 | - | - |
| Y = GAAB + IMD + [Volume] | 6 | 0.086 | <0.001 | 4.08 | 0.003 |
| Y = GAAB + IMD + [DTI] | 8 | 0.046 | 0.008 | 1.00 | 0.423 |
| Y = GAAB + IMD + [Volume] + [DTI] | 10 | 0.078 | <0.001 | 2.23 | 0.026 |
| **B. Motor** |  |  |  |  |  |
| **n=332** |  |  |  |  |  |
| Y = GAAB + IMD | 2 | 0.042 | 0.002 | - | - |
| Y = GAAB + IMD + [Volume] | 6 | 0.080 | <0.001 | 4.40 | 0.002 |
| **n=281** |  |  |  |  |  |
| Y = GAAB + IMD | 2 | 0.024 | 0.012 | - | - |
| Y = GAAB + IMD + [Volume] | 6 | 0.059 | 0.001 | 3.51 | 0.008 |
| Y = GAAB + IMD + [DTI] | 8 | 0.031 | 0.037 | 1.30 | 0.258 |
| Y = GAAB + IMD + [Volume] + [DTI] | 10 | 0.053 | 0.006 | 2.06 | 0.041 |
| **C. Communication** |  |  |  |  |  |
| **n=332** |  |  |  |  |  |
| Y = GAAB + IMD | 2 | 0.044 | 0.001 | - | - |
| Y = GAAB + IMD + [Volume] | 6 | 0.083 | <0.001 | 3.93 | 0.004 |
| **n=281** |  |  |  |  |  |
| Y = GAAB + IMD | 2 | 0.048 | <0.001 | - | - |
| Y = GAAB + IMD + [Volume] | 6 | 0.089 | <0.001 | 4.14 | 0.003 |
| Y = GAAB + IMD + [DTI] | 8 | 0.082 | <0.001 | 3.05 | 0.011 |
| Y = GAAB + IMD + [Volume] + [DTI] | 10 | 0.090 | <0.001 | 2.85 | 0.007 |

[Volume] = mean feature value (Jacobians) of regions positively associated with gestational age at birth, mean feature value of regions negatively associated with gestational age at birth, post menstrual age at scan, sex [DTI] = mean feature value of FA regions positively associated with gestational age at birth, mean feature value of MD, AD and RD regions negatively associated with gestational age at birth, post menstrual age at scan, sex.

## Discussion

Gestational age at birth related MRI phenotypes were observed in a cohort of healthy term-born babies. Lower gestational age at birth was also associated with poorer neurodevelopment, and individual differences in neurodevelopment may be partially explained by neonatal brain biology.

Infants born earlier had lower deep grey matter volumes, lower cerebellar volumes, and relatively larger ventricular volume (Figure 1). This anatomical pattern is similar to that observed in babies born late preterm (Gui et al., 2019). The pattern of white matter microstructural change seen here in infants born earlier (lower FA, higher MD, AD and RD) is also similar to that observed in late preterm infants compared to term controls (Kim et al., 2016).

In addition to the neuroimaging results, we demonstrate a GAAB association with neurodevelopmental outcome at 18 months, as assessed with the Bayley-III scales. This finding adds to a body of literature demonstrating poorer average performance on cognitive tests in individuals born in the early term period (Chan et al., 2016). Our results are broadly in keeping with one previous study which demonstrated a linear association between GAAB and Bayley-Il sub-scales at 12 months in a cohort of 232 infants from Southern California, reporting an r^2^ value of 0.12 for the Mental Development Index and 0.08 for the Psychomotor Development Index (Espel et al., 2014).

We build upon these studies by demonstrating how inter-individual brain differences may also affect later neurodevelopmental outcome. Two direct imaging – outcome associations were observed: the volume of brain regions which negatively correlated with GAAB also negatively correlated with Bayley-III motor (adj.r^2^ = 0.04, p<0.001) and cognitive score (adj.r^2^ = 0.02, p=0.02), although the latter of these would not survive multiple comparisons correction (Figure 4). These results are interesting but should be viewed with a degree of caution, the regions of interest used were of course defined by their association with GAAB in this specific cohort, which itself correlates with Bayley-III scores (Figure 3). Adding mean feature values of the relative volume (Jacobians) in these regions significantly improved linear regression models of GAAB-outcome, although adding mean feature values of white matter regions significantly associated with GAAB only improved the model of Communication (Table 2).

It is not clear why individual volume differences appear to be more important in future neurodevelopmental outcome than diffusion differences, although this is in keeping with existing research. Relative brain volume changes observed in the fetal or early neonatal period, especially ventricular enlargement (as seen in our cohort), have been linked to later neurodevelopmental outcome in several other cohorts (Alarcon et al., 2013; Hahner et al., 2019; Mirmiran et al., 2004), although the populations in these studies were either preterm or had an established clinical risk, which makes comparison to our cohort of healthy term-born babies difficult. This is a general trend in the literature; previously published studies relating neonatal MRI to neurodevelopment in term age babies use populations that have a pre-existing increased risk of a negative outcome, such as punctate lesions (Hayman et al., 2019), neonatal stroke (Dinomais et al., 2015), neonatal critical illness (Ramadan et al., 2012), or high parental age (Gale-Grant et al., 2020).

Our study has some limitations. The most significant of these is the mathematical difficulty of delineating the effect of GAAB on MRI phenotype with the much more dominant effect of brain maturation over time, as measured by post menstrual age at scan. In a cohort of term-born babies scanned postnatally these two variables are colinear (as is the case in our cohort, Supplementary Figure 1). Some of the areas whose volume positively correlates with GAAB also negatively correlate with gestational age at scan, and vice versa: for example the basal ganglia and brainstem, which may represent over correction of the effect of temporal maturation (Figure 1, Supplementary Figure 2). However, it is unlikely that over correction of brain maturation accounts for all of the GAAB associated brain volume differences seen here as the volumes of the ventricles (negatively associated with GAAB) were not associated with post menstrual age at scan, and the volume of parts of the cerebellum were positively associated with both GAAB and post menstrual age at scan. The areas of the white matter skeleton whose FA is significantly positively associated with GAAB are also significantly positively associated with post menstrual age at scan, and the areas of MD, AD and RD significantly negatively associated with GAAB are significantly negatively associated with post menstrual age at scan.

At the time of writing, all previously published studies which associated MRI findings in the neonatal period with GAAB (Broekman et al., 2014; Jin et al., 2019; Ou et al., 2017) use postnatal days of life as a covariate rather than pot menstrual age at scan. This likely gives results which are more representative of general maturation than of the effect of GAAB. This phenomenon can be illustrated by repeating our analysis of the association of GAAB and brain volume with postnatal age as a covariate in the place of post menstrual age at scan (Supplementary Figure 3). The results are very similar to the effect of post menstrual age (Supplementary Figure 2) but are not similar to the effect of GAAB (Figure 1).

Our study has several strengths. We have used a large homogenous cohort, which improves the quality of neuroimaging analysis and removes potential bias from abnormally low birthweights and medical complications in early term but does limit clinical translatability of our findings. We have used well established MRI analysis methods. An immediate avenue for future research would be to investigate the effect of GAAB on functional connectivity measures in the neonatal period as these have been shown to be significant in older children (Kim et al., 2014).

These results are important for future neuroimaging studies. Neonatal MRI is now well established within the neuroimaging field, and an increasing number of studies focus on term-born infants. Many of these do not include GAAB as a covariate in analyses (Feldmann et al., 2020; Glass et al., 2021; Merhar et al., 2020) – this study firmly suggests that analyses of MRI phenotype in term-born cohorts should be GAAB corrected. The results are also interesting from a clinical perspective. 37 weeks gestational age is widely viewed as a cut-off at which labour may be optimally induced if there is an indication to do so – 8% of all births in the USA in 2006 were induced in early term (Murthy et al., 2011). Caesarean section is also routinely electively performed before 40 weeks - guidelines in both the UK and the USA recommend any time from 39 weeks, although both cite perioperative morbidity rather than any concern about infant neurodevelopment as their rationale for this gestational age (The American College of Obstetricians and Gynecologists, 2019; The National Institute for Health and Care Excellence, 2011). Our results add to a body of literature demonstrating specific phenotypes associated with early term birth which merit further investigation.

This work advances our understanding of the neurodevelopmental effects of birth at different gestational ages after 37 weeks. The finding of significant associations of GAAB with brain phenotype soon after birth is important to consider in neonatal neuroimaging studies.

## Funding

This work was supported by the European Research Council under the European Union’s Seventh Framework Programme (FP7/20072013)/ERC grant agreement no. 319456 (dHCP project). The authors acknowledge infrastructure support from the National Institute for Health Research (NIHR) Mental Health Biomedical Research Centre (BRC) at South London, Maudsley NHS Foundation Trust, King’s College London and the NIHR-BRC at Guys and St Thomas’ Hospitals NHS Foundation Trust (GSTFT). The authors also acknowledge support in part from the Wellcome Engineering and Physical Sciences Research Council (EPSRC) Centre for Medical Engineering at Kings College London [WT 203148/Z/16/Z], MRC strategic grant [MR/K006355/1], Medical Research Council Centre grant [MR/N026063/1], the Department of Health through an NIHR Comprehensive Biomedical Research Centre Award (to Guy’s and St. Thomas’ National Health Service (NHS) Foundation Trust in partnership with King’s College London and King’s College Hospital NHS Foundation Trust), the Sackler Institute for Translational Neurodevelopment at King’s College London and the European Autism Interventions (EU-AIMS) trial and the EU AIMS-2-TRIALS, a European Innovative Medicines Initiative Joint Undertaking under Grant Agreements No. 115300 and 777394, the resources of which are composed of financial contributions from the European Union’s Seventh Framework Programme (Grant FP7/2007–2013). OGG is supported by a grant from the UK Medical Research Council [MR/P502108/1]. JOM and DE received support from the Medical Research Council Centre for Neurodevelopmental Disorders, King’s College London [MR/N026063/1]. LCG is supported by a Beatriz Galindo Fellowship jointly funded by the Ministerio de Educación, Cultura y Deporte and the Universidad Politécnica de Madrid [BEAGAL18/00158]. JOM is supported by a Sir Henry Dale Fellowship jointly funded by the Wellcome Trust and the Royal Society [206675/Z/17/Z]. TA is supported by a MRC Clinician Scientist Fellowship [MR/P008712/1] and Transition Support Award [MR/V036874/1].DB received support from a Wellcome Trust Seed Award in Science [217316/Z/19/Z]. The views expressed are those of the authors and not necessarily those of the NHS, the National Institute for Health Research or the Department of Health. The funders had no role in the design and conduct of the study; collection, management, analysis, and interpretation of the data; preparation, review, or approval of the manuscript; and decision to submit the manuscript for publication.

**Supplementary Figure 1 –.**
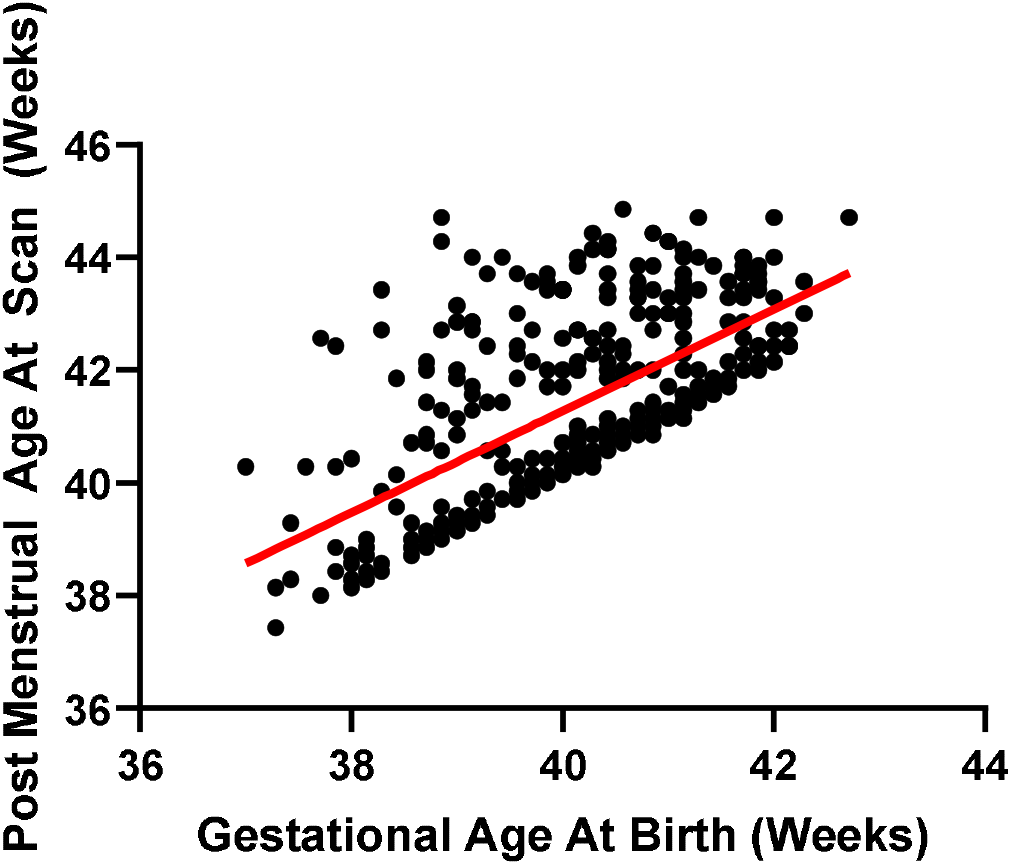
Association of Gestational Age at Birth with Post Menstrual Age at Scan. r^2^ =0.40, p<0.0001

**Supplementary Figure 2 –.**
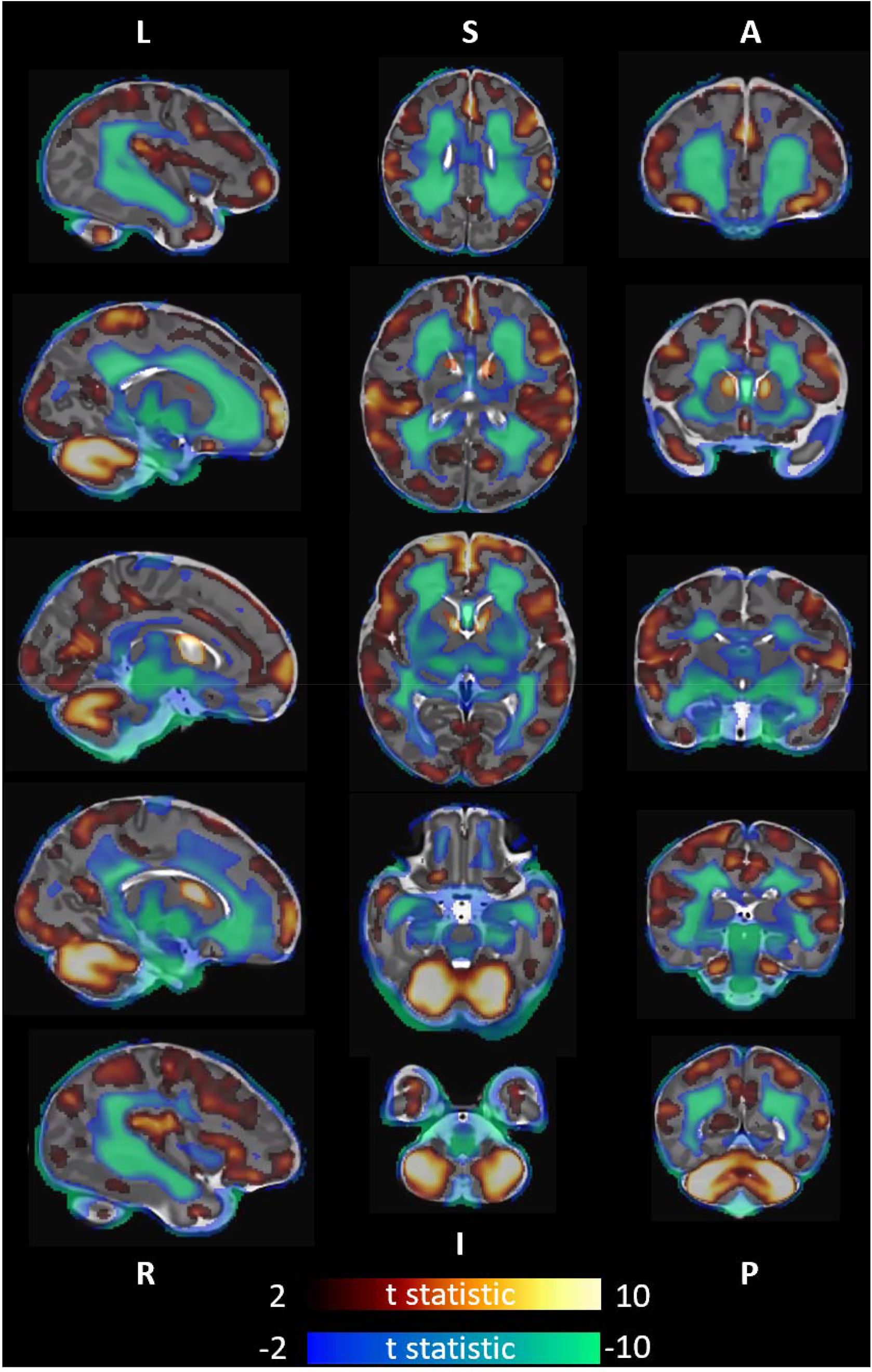
Association of post menstrual age at scan with brain volume. T statistic of areas of significant (p<0.025) correlation shown - positive in red-yellow, negative in blue-green. 1^st^ column left to right, 2^nd^ column superior to inferior, 3^rd^ column anterior to posterior. Analysis corrected for sex.

**Supplementary Figure 3 –.**
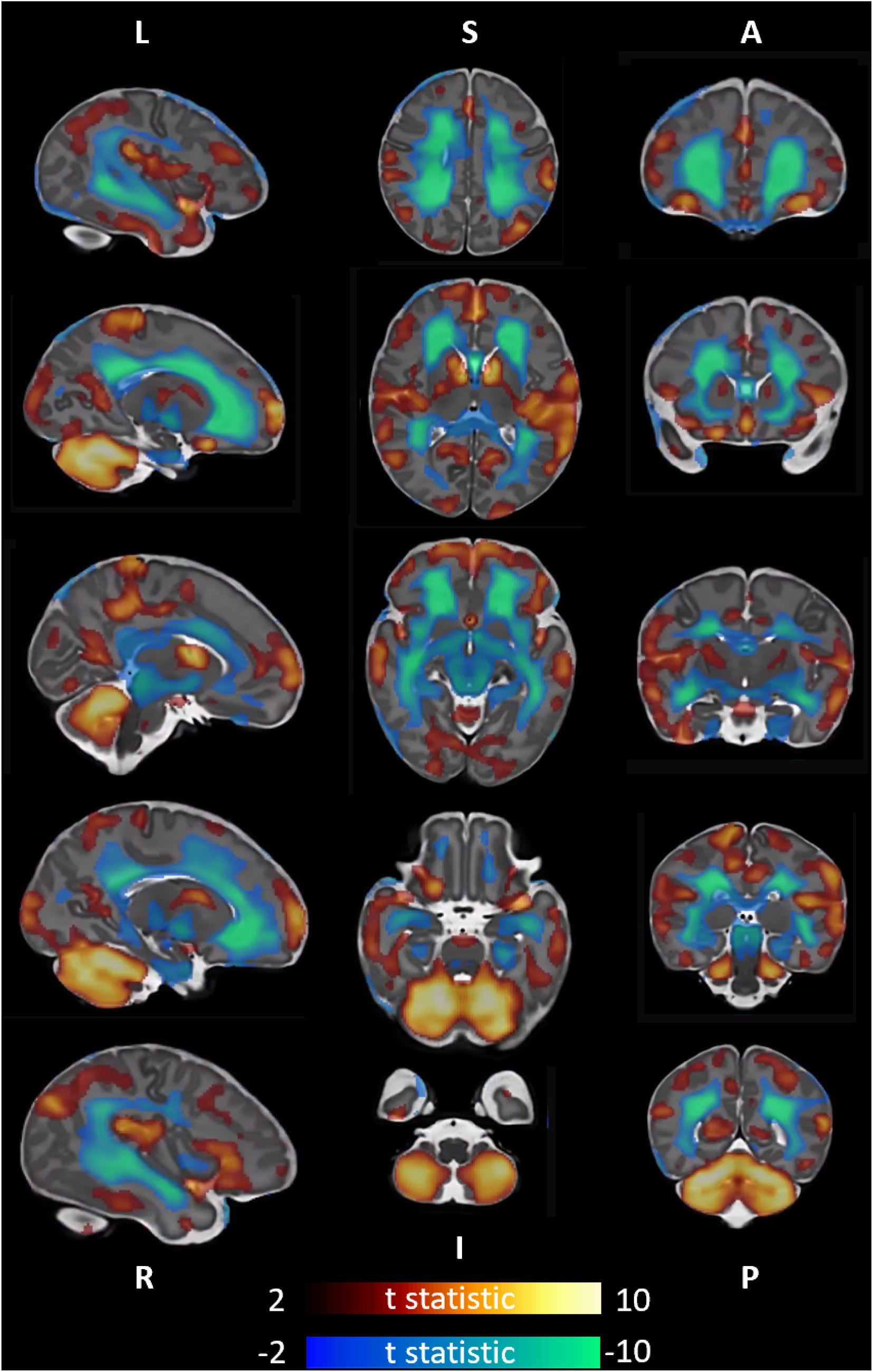
Association of gestational age at birth with brain volume, uncorrected for post menstrual age at scan. T statistic of areas of significant (p<0.025) correlation shown - positive in red-yellow, negative in blue-green. 1^st^ column left to right, 2^nd^ column superior to inferior, 3^rd^ column anterior to posterior. Analysis corrected for postnatal age (days) and sex.

## References

Abel, G.A., Barclay, M.E., Payne, R.A., 2016. Adjusted indices of multiple deprivation to enable comparisons within and between constituent countries of the UK including an illustration using mortality rates. BMJ Open 6, e012750.

Alarcon, A., Martinez-Biarge, M., Cabañas, F., Hernanz, A., Quero, J., Garcia-Alix, A., 2013. Clinical, biochemical, and neuroimaging findings predict long-term neurodevelopmental outcome in symptomatic congenital cytomegalovirus infection. J Pediatr 163, 828–834.e821.

Ashburner, J., Friston, K.J., 2000. Voxel-based morphometry--the methods. Neuroimage 11, 805–821.

Avants, B., Gee, J.C., 2004. Geodesic estimation for large deformation anatomical shape averaging and interpolation. Neuroimage 23 Suppl 1, S139–150.

Ball, G., Counsell, S.J., Anjari, M., Merchant, N., Arichi, T., Doria, V., Rutherford, M.A., Edwards, A.D., Rueckert, D., Boardman, J.P., 2010. An optimised tract-based spatial statistics protocol for neonates: applications to prematurity and chronic lung disease. Neuroimage 53, 94–102.

Batalle, D., Edwards, A.D., O’Muircheartaigh, J., 2018. Annual Research Review: Not just a small adult brain: understanding later neurodevelopment through imaging the neonatal brain. J Child Psychol Psychiatry 59, 350–371.

Bayley, N., 2006. Bayley Scales of Infant and Toddler Development-3rd Edition (Bayley-III). The Psychological Corporation, San Antonio, TX.

Boyle, E.M., Poulsen, G., Field, D.J., Kurinczuk, J.J., Wolke, D., Alfirevic, Z., Quigley, M.A., 2012. Effects of gestational age at birth on health outcomes at 3 and 5 years of age: population based cohort study. Bmj 344, e896.

Broekman, B.F., Wang, C., Li, Y., Rifkin-Graboi, A., Saw, S.M., Chong, Y.S., Kwek, K., Gluckman, P.D., Fortier, M.V., Meaney, M.J., Qiu, A., 2014. Gestational age and neonatal brain microstructure in term born infants: a birth cohort study. PLoS One 9, e115229.

Budin, P., 1907. The Nursling; The Feeding and Hygiene of Premature and Full-Term Infants. Caxton, London.

Chan, E., Leong, P., Malouf, R., Quigley, M.A., 2016. Long-term cognitive and school outcomes of late-preterm and early-term births: a systematic review. Child Care Health Dev 42, 297–312.

Christiaens, D., Cordero-Grande, L., Pietsch, M., Hutter, J., Price, A.N., Hughes, E.J., Vecchiato, K., Deprez, M., Edwards, A.D., Hajnal, J.V., Tournier, J.D., 2021. Scattered slice SHARD reconstruction for motion correction in multi-shell diffusion MRI. Neuroimage 225, 117437.

Coates, D., Makris, A., Catling, C., Henry, A., Scarf, V., Watts, N., Fox, D., Thirukumar, P., Wong, V., Russell, H., Homer, C., 2020. A systematic scoping review of clinical indications for induction of labour. PLoS One 15, e0228196.

Coathup, V., Boyle, E., Carson, C., Johnson, S., Kurinzcuk, J.J., Macfarlane, A., Petrou, S., Rivero-Arias, O., Quigley, M.A., 2020. Gestational age and hospital admissions during childhood: population based, record linkage study in England (TIGAR study). Bmj 371, m4075.

Cordero-Grande, L., Hughes, E.J., Hutter, J., Price, A.N., Hajnal, J.V., 2018. Three-dimensional motion corrected sensitivity encoding reconstruction for multi-shot multi-slice MRI: Application to neonatal brain imaging. Magn Reson Med 79, 1365–1376.

Davis, E.P., Sandman, C.A., 2010. The timing of prenatal exposure to maternal cortisol and psychosocial stress is associated with human infant cognitive development. Child Dev 81, 131–148.

Dinomais, M., Hertz-Pannier, L., Groeschel, S., Chabrier, S., Delion, M., Husson, B., Kossorotoff, M., Renaud, C., Nguyen The Tich, S., 2015. Long term motor function after neonatal stroke: Lesion localization above all. Hum Brain Mapp 36, 4793–4807.

Doctor, B.A., O’Riordan, M.A., Kirchner, H.L., Shah, D., Hack, M., 2001. Perinatal correlates and neonatal outcomes of small for gestational age infants born at term gestation. Am J Obstet Gynecol 185, 652–659.

El Marroun, H., Zou, R., Leeuwenburg, M.F., Steegers, E.A.P., Reiss, I.K.M., Muetzel, R.L., Kushner, S.A., Tiemeier, H., 2020. Association of Gestational Age at Birth With Brain Morphometry. JAMA Pediatr.

Espel, E.V., Glynn, L.M., Sandman, C.A., Davis, E.P., 2014. Longer gestation among children born full term influences cognitive and motor development. PLoS One 9, e113758–e113758.

Feldmann, M., Guo, T., Miller, S.P., Knirsch, W., Kottke, R., Hagmann, C., Latal, B., Jakab, A., 2020. Delayed maturation of the structural brain connectome in neonates with congenital heart disease. Brain Commun 2, fcaa209.

Gale-Grant, O., Christiaens, D., Cordero-Grande, L., Chew, A., Falconer, S., Makropoulos, A., Harper, N., Price, A.N., Hutter, J., Hughes, E., Victor, S., Counsell, S.J., Rueckert, D., Hajnal, J.V., Edwards, A.D., O’Muircheartaigh, J., Batalle, D., 2020. Parental age effects on neonatal white matter development. Neuroimage Clin 27, 102283.

Glass, H.C., Li, Y., Gardner, M., Barkovich, A.J., Novak, I., McCulloch, C.E., Rogers, E.E., 2021. Early Identification of Cerebral Palsy Using Neonatal MRI and General Movements Assessment in a Cohort of High-Risk Term Neonates. Pediatr Neurol 118, 20–25.

Gui, L., Loukas, S., Lazeyras, F., Hüppi, P.S., Meskaldji, D.E., Borradori Tolsa, C., 2019. Longitudinal study of neonatal brain tissue volumes in preterm infants and their ability to predict neurodevelopmental outcome. Neuroimage 185, 728–741.

Hahner, N., Benkarim, O.M., Aertsen, M., Perez-Cruz, M., Piella, G., Sanroma, G., Bargallo, N., Deprest, J., Gonzalez Ballester, M.A., Gratacos, E., Eixarch, E., 2019. Global and Regional Changes in Cortical Development Assessed by MRI in Fetuses with Isolated Nonsevere Ventriculomegaly Correlate with Neonatal Neurobehavior. AJNR Am J Neuroradiol 40, 1567–1574.

Hayman, M., van Wezel-Meijler, G., van Straaten, H., Brilstra, E., Groenendaal, F., de Vries, L.S., 2019. Punctate white-matter lesions in the full-term newborn: Underlying aetiology and outcome. Eur J Paediatr Neurol 23, 280–287.

Hua, J., Sun, J., Cao, Z., Dai, X., Lin, S., Guo, J., Gu, G., Du, W., 2019. Differentiating the cognitive development of early-term births in infants and toddlers: a cross-sectional study in China. BMJ Open 9, e025275.

Hughes, E.J., Winchman, T., Padormo, F., Teixeira, R., Wurie, J., Sharma, M., Fox, M., Hutter, J., Cordero-Grande, L., Price, A.N., Allsop, J., Bueno-Conde, J., Tusor, N., Arichi, T., Edwards, A.D., Rutherford, M.A., Counsell, S.J., Hajnal, J.V., 2017. A dedicated neonatal brain imaging system. Magn Reson Med 78, 794–804.

Hutter, J., Tournier, J.D., Price, A.N., Cordero-Grande, L., Hughes, E.J., Malik, S., Steinweg, J., Bastiani, M., Sotiropoulos, S.N., Jbabdi, S., Andersson, J., Edwards, A.D., Hajnal, J.V., 2018. Time-efficient and flexible design of optimized multishell HARDI diffusion. Magn Reson Med 79, 1276–1292.

Jenkinson, M., Beckmann, C.F., Behrens, T.E., Woolrich, M.W., Smith, S.M., 2012. FSL. Neuroimage 62, 782–790.

Jin, C., Li, Y., Li, X., Liu, C., Wang, M., Cheng, Y., Zheng, J., Yang, J., 2019. Associations of gestational age and birth anthropometric indicators with brain white matter maturation in full-term neonates. Hum Brain Mapp 40, 3620–3630.

Kellner, E., Dhital, B., Kiselev, V.G., Reisert, M., 2016. Gibbs-ringing artifact removal based on local subvoxel-shifts. Magn Reson Med 76, 1574–1581.

Kelly, C.J., Christiaens, D., Batalle, D., Makropoulos, A., Cordero-Grande, L., Steinweg, J.K., O’Muircheartaigh, J., Khan, H., Lee, G., Victor, S., Alexander, D.C., Zhang, H., Simpson, J., Hajnal, J.V., Edwards, A.D., Rutherford, M.A., Counsell, S.J., 2019. Abnormal Microstructural Development of the Cerebral Cortex in Neonates With Congenital Heart Disease Is Associated With Impaired Cerebral Oxygen Delivery. J Am Heart Assoc 8, e009893.

Kim, D.-Y., Park, H.-K., Kim, N.-S., Hwang, S.-J., Lee, H.J., 2016. Neonatal diffusion tensor brain imaging predicts later motor outcome in preterm neonates with white matter abnormalities. Italian journal of pediatrics 42, 104–104.

Kim, D.J., Davis, E.P., Sandman, C.A., Sporns, O., O’Donnell, B.F., Buss, C., Hetrick, W.P., 2014. Longer gestation is associated with more efficient brain networks in preadolescent children. Neuroimage 100, 619–627.

Malamitsi-Puchner, A., 2017. Preterm birth in ancient Greece: a synopsis. J Matern Fetal Neonatal Med 30, 141–143.

Merhar, S.L., Kline, J.E., Braimah, A., Kline-Fath, B.M., Tkach, J.A., Altaye, M., He, L., Parikh, N.A., 2020. Prenatal opioid exposure is associated with smaller brain volumes in multiple regions. Pediatr Res.

Mirmiran, M., Barnes, P.D., Keller, K., Constantinou, J.C., Fleisher, B.E., Hintz, S.R., Ariagno, R.L., 2004. Neonatal brain magnetic resonance imaging before discharge is better than serial cranial ultrasound in predicting cerebral palsy in very low birth weight preterm infants. Pediatrics 114, 992–998.

Murthy, K., Grobman, W.A., Lee, T.A., Holl, J.L., 2011. Trends in induction of labor at early-term gestation. Am J Obstet Gynecol 204, 435.e431–436.

Murzakanova, G., Räisänen, S., Jacobsen, A.F., Sole, K.B., Bjarkø, L., Laine, K., 2020. Adverse perinatal outcomes in 665,244 term and post-term deliveries-a Norwegian population-based study. Eur J Obstet Gynecol Reprod Biol 247, 212–218.

Ou, X., Glasier, C.M., Ramakrishnaiah, R.H., Kanfi, A., Rowell, A.C., Pivik, R.T., Andres, A., Cleves, M.A., Badger, T.M., 2017. Gestational Age at Birth and Brain White Matter Development in Term-Born Infants and Children. AJNR Am J Neuroradiol 38, 2373–2379.

Ramadan, G., Paul, N., Morton, M., Peacock, J.L., Greenough, A., 2012. Outcome of ventilated infants born at term without major congenital abnormalities. Eur J Pediatr 171, 331–336.

Rose, O., Blanco, E., Martinez, S.M., Sim, E.K., Castillo, M., Lozoff, B., Vaucher, Y.E., Gahagan, S., 2013. Developmental scores at 1 year with increasing gestational age, 37-41 weeks. Pediatrics 131, e1475–1481.

Schuh, A., Makropoulos, A., Robinson, E.C., Cordero-Grande, L., Hughes, E., Hutter, J., Price, A.N., Murgasova, M., Teixeira, R.P.A.G., Tusor, N., Steinweg, J.K., Victor, S., Rutherford, M.A., Hajnal, J.V., Edwards, A.D., Rueckert, D., 2018. Unbiased construction of a temporally consistent morphological atlas of neonatal brain development. bioRxiv, 251512.

Taylor, H.G., Minich, N., Bangert, B., Filipek, P.A., Hack, M., 2004. Long-term neuropsychological outcomes of very low birth weight: associations with early risks for periventricular brain insults. J Int Neuropsychol Soc 10, 987–1004.

The American College of Obstetricians and Gynecologists, 2019. ACOG Committee Opinion No. 761: Cesarean Delivery on Maternal Request. Obstet Gynecol 133, e73–e77.

The National Institute for Health and Care Excellence, 2011. Clinical guideline [CG132]: Caesarean section. In: NICE (Ed.). NICE, London.

Winkler, A.M., Ridgway, G.R., Webster, M.A., Smith, S.M., Nichols, T.E., 2014. Permutation inference for the general linear model. Neuroimage 92, 381–397.

Zhang, H., Yushkevich, P.A., Alexander, D.C., Gee, J.C., 2006. Deformable registration of diffusion tensor MR images with explicit orientation optimization. Medical Image Analysis 10, 764–785.

